# Proteome dynamics during antibiotic persistence and resuscitation

**DOI:** 10.1101/2020.09.04.282723

**Authors:** Maja Semanjski, Fabio Gratani, Till Englert, Payal Nashier, Viktor Beke, Nicolas Nalpas, Elsa Germain, Shilpa George, Christiane Wolz, Kenn Gerdes, Boris Macek

## Abstract

During antibiotic persistence, bacterial cells become transiently tolerant to antibiotics by restraining their growth and metabolic activity. Detailed molecular characterization of antibiotic persistence is hindered by low count of persisting cells and the need for their isolation. Here we used sustained addition of stable isotope-labeled lysine to selectively label the proteome during *hipA*-induced persistence and *hipB*-induced resuscitation of *E. coli* cells in minimal medium after antibiotic treatment. Time-resolved, 24-hour measurement of label incorporation allowed detection of over 500 newly synthesized proteins in viable cells, demonstrating low but widespread protein synthesis during persistance. Many essential proteins were newly synthesized and several ribosome-associated proteins, such as RaiA and Sra showed high synthesis levels, pointing to their roles in maintenance of persistence. At the onset of resuscitation, cells synthesized the ribosome-splitting GTPase HflX and various ABC transporters, restored translation machinery and resumed metabolism by inducing glycolysis and biosynthesis of amino acids.

## Introduction

Antibiotic resistance is an acute health problem. Many bacteria are categorized as serious threats representing a considerable clinical and financial burden [1, 2]. In addition to resistance, bacteria also possess an elusive innate strategy, termed persistence, that enables them to tolerate antibiotics without acquiring genetic changes [3]. Persisters are defined as phenotypic variants of bacterial cells that become transiently tolerant to antibiotics by restraining their growth and entering a dormant-like state [4, 5]. Persistence is exhibited within a subpopulation of genetically uniform cells, a phenomenon present in all bacteria tested so far [6, 7]. While bactericidal antibiotics typically require actively growing cells to exploit their function, persister cells are slowly replicating which makes them tolerant to the lethal action of antimicrobials. In addition to slow growth, the persister phenotype is also associated with very low metabolic activity [8].

Persistence can be induced by toxin-antitoxin (TA) modules that usually consist of two genes: one encoding a toxin that inhibits cell growth, and another encoding an antitoxin that inhibits toxin activity [9, 10]. While the product of the toxin gene is a protein that interferes with essential cellular functions, antitoxin genes encode either small proteins or noncoding RNAs [9]. Several studies have shown that persister cultures are heterogenous, with large abundance of cells with intact membranes that are viable but not culturable (VBNC) [11–14]. In addition, low counts of persister cells in these cultures necessitate application of elaborate protocols for their isolation [15–17], making detailed, unbiased exploration of the unperturbed molecular processes a challenging task.

Here we used an *in vitro* model of bacterial persistence based on conditional overproduction of HipA toxin and HipB antitoxin [18–23] to devise a generic method for temporal analysis of protein synthesis during toxin-induced persistence and antitoxin-mediated resuscitation. We reasoned that prolonged antibiotic treatment of HipA-induced *E. coli* cells, which results in a mixture of dead (lysed), and viable cells, can be coupled with sustained addition of stable isotope-labeled amino acid directly to the culture. As only the viable, persisting cells are capable of incorporating the label, and the label mass shift can be easily resolved in a mass spectrometer, this approach can be used to selectively label and detect newly synthesized proteins in the high background of proteins originating from dead cells. For simplicity, in this manuscript we will use the term “persisters” for all cells that remain viable after this treatment; however, we note that this population is heterogenous and consists of culturable and viable but non-culturable (VBNC) cells. Similar strategies, termed “pulsed” or “dynamic” stable isotope labeling by amino acids in cell culture (SILAC), have previously been used to analyze protein turnover in eukaryotic [24, 25] and prokaryotic cells [26, 27].

## Results

### HipA-induced persister cells show low but measurable protein synthesis

To test applicability of our approach, we first increased the number of persister cells growing in minimal medium by ectopically inducing *hipA* in *E. coli* MG1655. Three hours after HipA-induced growth inhibition, we treated the culture with ampicillin to lyse antibiotic-sensitive cells. During ampicillin treatment, we repressed expression of *hipA* by addition of glucose to ensure steady state conditions independent of the constant HipA overproduction. Twenty hours after ampicillin treatment, we added ^15^N2^13^C6-lysine (Lys8) to the culture; this time point was sufficient to eradicate sensitive cells and achieve a steady state of persistence, as demonstrated by approximately 1000-fold higher count of colony-forming units (CFUs) in *hipA*-induced strain compared to the empty vector (**Supplementary Figure 1A,B**). After introduction of the Lys8 label, we harvested the culture aliquots by brief centrifugation to pellet intact cells in three biological replicates at multiple time points ranging from 10 minutes to 24 hours (**Figure 1A**). In this experimental setup, only viable cells incorporated Lys8, which enabled specific detection of newly synthesized proteins, temporal quantification of protein abundance and estimation of protein half-lives (**Supplementary Figure 1C, Supplementary Table S1**). The number of identified proteins across time points was consistent, with 1,993 proteins identified on average in each time point (**Supplementary Figure 1D**). Remarkably, up to 531 proteins have partially incorporated the Lys8 label and were reproducibly quantifiable in the high background of Lys0-labeled proteins originating from dead cells (**Supplementary Figure 1D,E**). Due to HipA-mediated inhibition of cell growth, incorporation of Lys8 in these proteins was only partial and expectedly low; 24 hours after the start of labeling the average intensity of all heavy labeled signals amounted to only 8.64% of the total ion intensity (**Figure 1B**). Nevertheless, the measurable label incorporation was indicative of protein synthesis in viable cells and was in agreement with previous studies [27–29]. The low level of label incorporation reflected the low turnover rates and long half-lives of proteins in persisting cells. Estimation of protein half-lives from the measured H/L ratios, as described previously [24], revealed that the median protein half-life under these conditions was longer than 250 hours (**Supplementary Figure 1F, Supplementary Table S1**). This, however, is only an estimate, since our measurement window (24h) was significantly shorter than the actual protein half-life and our calculation was based on the assumption of complete cell cycle arrest. This experiment demonstrated that it is possible to partially label and directly detect newly synthesized proteins of viable cells after toxin overproduction and antibiotic treatment. All measured proteins and their ratios are listed in the **Supplementary Table S2** and can be browsed online under https://pctsee.pct.uni-tuebingen.de.

**Figure 1.**
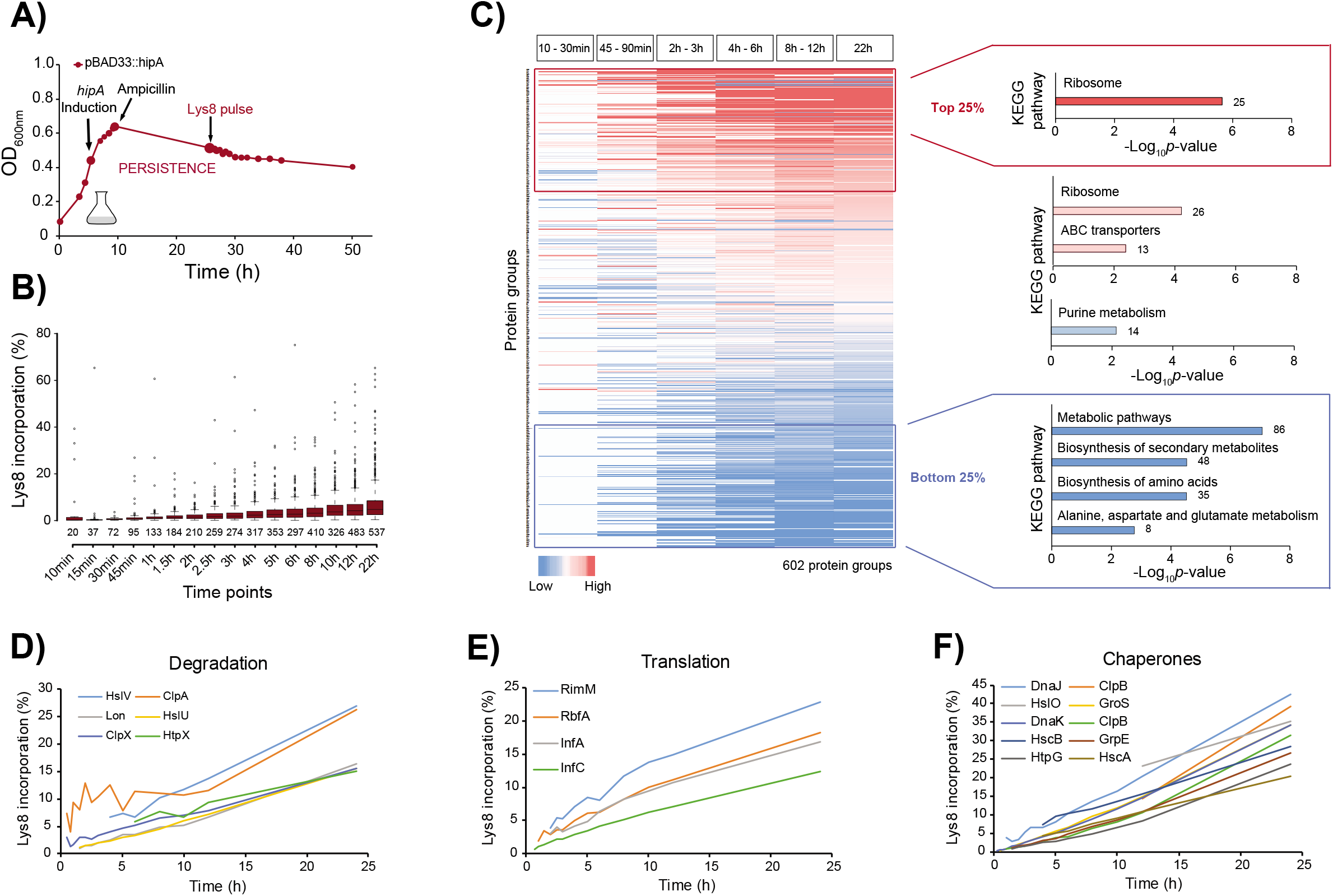
Persister cells synthesize proteins involved in essential physiological processes. **(A)** Transcription of plasmid-encoded *hipA* (pBAD33::*hipA*) was induced in *E. coli* K-12 strain MG1655 at an optical density OD_600nm_ of 0.4 for 3 hours, followed by ampicillin treatment and addition of glucose to repress further *hipA* transcription. Approx. 20 hours after *hipA* induction, the culture was sampled at 16 time points. Proteins were digested with endoproteinase Lys-C and measured by LC-MS/MS. The growth curve is representative of three biological replicates. **(B)** Incorporation of Lys8 into proteins across 17 time points calculated from the measured protein H/L ratios. The box plots are representative of 3 biological replicates. **(C)** Heat map of all quantified proteins color-coded based on their H/L ratio ranking within each time bin. Missing values are colored in white. KEGG pathway enrichment analysis of quanified proteins in the 25th percentile (Top 25%, red), 25th – 50th percentile (pink), 50th – 75th percentile (light red and blue) and 75th percentile (Bottom 25%, blue) across all time bins. The number of enriched proteins in each category is indicated outside the bars. **(D-F)** Label incorporation curves of selected examples of enzymes involved in protein degradation, translation and folding (chaperones).

### Persisting bacteria produce proteins needed for essential cellular processes

For further data analysis, the time points over the entire 24-hour course of persistence were collapsed into six time bins and proteins were ranked based on their H/L ratios. Gene Ontology (GO) analysis of all newly synthesized proteins showed that proteins involved in translation, glycolysis, protein targeting, redox homeostasis and tricarboxylic-acid cycle were significantly enriched (*p* < 0.01) (**Supplementary Table S3**). Importantly, almost all ribosomal proteins were overrepresented among top 25% proteins that exhibited relatively high label incorporation across all time points (*p* = 1.67E-15). In contrast, proteins with lower label incorporation (bottom 25%) participate mainly in the metabolic pathways such as biosynthesis of amino acids and other metabolites (**Figure 1C**). Interestingly, enzymes involved in protein degradation, including degradation of antitoxins necessary to maintain an optimal toxin/antitoxin ratio during persistence, also exhibited higher label incorporation. They included several proteases (Lon, HslU, HslV and HtpX), as well as the specificity components of the Clp protease (ClpA and ClpX, but not ClpP), implying that protein degradation is active during persistence (**Figure 1D**). Two antitoxins, PrlF and MqsA, as well as several heat and cold shock proteins involved in stress response, also showed Lys8 incorporation. Furthermore, proteins involved in transcription, such as the ribonucleases RNase R (Rnr) and RNase III (Rnc) required for RNA processing and turnover, including the RNA polymerase sigma factor RpoD, were also detected as newly synthesized, as were several translation regulatory proteins essential for the start of protein synthesis, such as translation initiation factors IF-1 and 1F-3 (InfA and InfC) (**Figure 1E**). Several chaperones were detected as newly synthesized during persistence (**Figure 1F**), suggesting that they may be required for the maintenance of proteome homeostasis, either by enabling correct folding of misfolded client proteins or by targeting unfolded (or aggregated) proteins for degradation. Among them were DnaK, DnaJ and ClpB, that were previously connected to persistence [30, 31]. Moreover, one of the most important DNA repair proteins RecA was identified as newly synthesized in our study, which is in agreement with a previous report that SOS response genes are expressed during persistence due to inhibited DNA replication [31]. These results indicate that bacterial cells perform active translation during antibiotic persistence and produce proteins that are needed for essential cellular processes. In fact, out of 299 essential gene products in *E. coli*, 159 incorporated the label during persistence (**Supplementary Table S4**).

### Ribosome-associated proteins RaiA and Sra show elevated synthesis levels during persistence

Consistent with the slow growth attributed to persister cells, proteins required for cell division (FtsH, FtsZ, FtsY and ZipA) were either not detected or displayed very low label incorporation in our data set, suggesting that residual cell growth is still present, but is extremely slow. Accordingly, chemotaxis-related proteins that are important for cell motility were not identified, which supports the absence of the cell movement during persistence. The low abundance of newly synthesized proteins involved in metabolism, mainly in biosynthesis of amino acids and primary carbon metabolism, demonstrated that major energy-generating pathways are active in our model of persistence, but their activity is very low. In contrast to reduced metabolism and cell division, we found that ribosomal proteins and general stress response-related proteins were incorporating the Lys8 label to a higher extent during persistence.

Regulation of ribosome function is known to have a key role in bacterial persistence [32]; therefore, we next analyzed relative abundance and label incorporation of all detected ribosomal proteins, as well as ribosome-associated hibernation and stationary phase factors proposed to modulate ribosomal activity, such as EttA, RaiA, ElaB, YqjD and Sra [32–34]. In persisting cells, measured rates of label incorporation and relative abundances of most ribosomal proteins were relatively high (**Figure 2A**). Of hibernation factors, only RaiA was detected as newly synthesized at a relatively high rate, pointing to its potential role as a major factor in active maintenance of ribosome hibernation during persistence. Intriguingly, the stationary phase ribosome associated factor Sra showed an even higher label incorporation rate and is therefore likely to be involved in the maintenance of ribosome hibernation (**Figure 2A**). Importantly, the level of newly synthesized GTPase HflX, proposed to mediate activation of hibernating ribosome dimers [35], was not detectable during persistence. Strikingly, the overall abundance of newly synthesized hibernation factors (predominantly RaiA), estimated using intensity-based absolute quantification (iBAQ) of the heavy isotope channel, were about 100 times higher than the level of all newly synthesized ribosomal proteins combined (**Figure 2B, Supplementary Table S5**).

**Figure 2.**
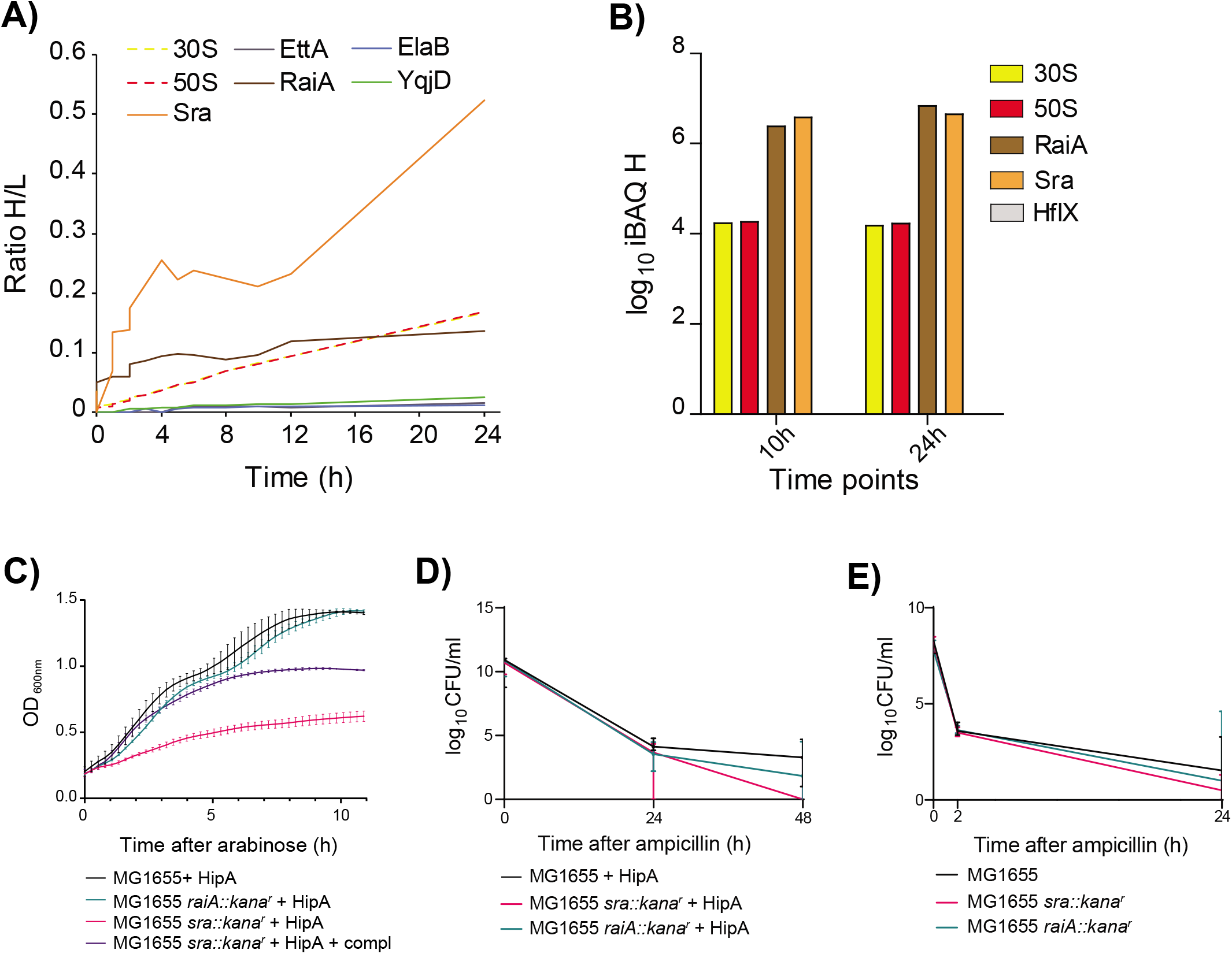
Ribosome-associated proteins RaiA and Sra show elevated synthesis levels in persisting cells and a role during antibiotic tolerance. **(A)** All detected ribosomal proteins had comparable label incorporation rates. The synthesis rate of the hibernation factor RaiA was elevated and similar to ribosomal proteins, whereas the synthesis rate of the ribosome-associated protein Sra was markedly higher than that of the ribosomal proteins. Most of the ribosome-associated hibernation factors (e.g. EttA, ElaB, YqjD) showed very low synthesis levels. **(B)** Estimates of the amounts of newly synthesized proteins based on intensity-based absolute quantification (iBAQ) [24] in the heavy SILAC channel. The amount of newly synthesized RaiA and Sra proteins was about 100 times higher than all ribosomal proteins combined. Depicted are the median values of three biological replicates. **(C)** WT and deletion strains *ΔraiA* presented similar growth phenotype upon HipA ectopic expression while strain *Δsra* deletion showed a stronger growth inhibition upon HipA expression, which can be partially complemented by plasmid bearing *sra* gene under tis native promoter. Strains bearing with empty vectors are used as controls. **(D)** WT strains ectopically expressing HipA were more viable compared to the same strain bearing the empty vector. Two deletion strains (*Δsra* and *ΔraiA*) expressing *hipA* showed a strong decrease in CFU, indicating potential role of Sra and RaiA in the antibiotic-tolerance mechanism (results represent six different replicates from two different experiments in the LB medium). Equivalent experiments in the M9 medium are shown in Supplementary Figure 1H. Control experiments including expression of empty vector are presented in Supplementary Figure 1I. **(E)** Without HipA ectopic overexpression, only mutant lacking the *sra* gene showed a lower survival in the LB medium.

### Ribosome-associated protein Sra is involved in antibiotic tolerance

To analyse potential involvement of Sra and RaiA in the HipA-induced growth inhibition, we expressed *hipA* in the *sra* and *raiA* deletion mutants at the beginning of the exponential phase and followed their growth over time (**Figure 2C, Supplementary figure 1G**). While *raiA* mutant showed a comparable growth curve to the WT upon *hipA* expression (**Figure 2C**), *sra* mutant showed a stronger inhibition, which could be partially complemented with a plasmid bearing *sra* gene under its native promoter (**Figure 2C**). To further test whether ribosome-associated proteins Sra and RaiA influence HipA-induced persistence and antibiotic tolerance, we ectopically induced *hipA* in *E. coli sra* and/or *raiA* deletion strains in the log phase and treated the cultures grown in LB medium with ampicillin for up to 48 hours. Already 24 hours after addition of antibiotic, the CFU counts of the *hipA*-expressing *sra* and *raiA* deletion mutants were lower compared to the *hipA*-expressing WT, pointing to a lower tolerance of these strains to ampicillin (**Figure 2D, Supplementary figure 1H**). A similar effect was also observed in the M9 medium (**Supplementary Figure 1I**). The difference in the CFU count was even more pronounced 48 hours after addition of antibiotic, with the CFU count of *raiA* mutant about 1E2-fold lower, and *sra* mutant 1E3-fold lower than in the *hipA*-expressing WT. A less pronounced but similar trend was also observed when *hipA* was not expressed (**Figure 2E**). These experiments strongly suggested involvement of ribosome-associated proteins, especially Sra, in maintenance of antibiotic tolerance.

### HipB-induction leads to rapid resuscitation and incorporation of the Lys8 label

We next applied our approach to study proteins produced upon induction of resuscitation. As in the first experiment, we increased the number of persister cells by ectopically inducing *hipA* in minimal medium supplemented with Lys0 and treated the culture with a high dose of ampicillin. During ampicillin treatment, we repressed expression of *hipA* by addition of glucose to ensure steady state conditions independent of any HipA overproduction. To trigger resuscitation, we removed ampicillin, Lys0 and cellular extract of lysed cells by gentle filtration and added fresh minimal medium containing IPTG to induce transcription of antitoxin *hipB*. The medium also contained “heavy” lysine (Lys8) for metabolic labeling of resuscitating cells. Filtered cultures exposed to the fresh, IPTG-containing medium resumed growth, whereas unfiltered control cultures did not (**Supplementary Figure 2A**), demonstrating that resuscitation was driven by *hipB* induction. The resuscitating culture was harvested in three biological replicates and 20 time points (**Figure 3A**). Incorporation of Lys8 in resuscitating cells followed the shape of the growth curve and saturated at average 94.6% (**Figure 3B**). Already 30 minutes after induction of resuscitation, more than 100 proteins incorporated the Lys8 label and could be quantified. Although the median of Lys8 incorporation during the first four hours was low (< 8%), presence of outliers indicated that a subset of proteins incorporated the label faster than others and are therefore likely to play a role in resuscitation. On average, 1,880 proteins were identified at an estimated FDR of 1.3% (**Supplementary Figure 2B**). The three biological replicates showed good reproducibility of measured protein SILAC ratios across all time points (**Supplementary Figure 2C; Supplementary Table S6**).

**Figure 3.**
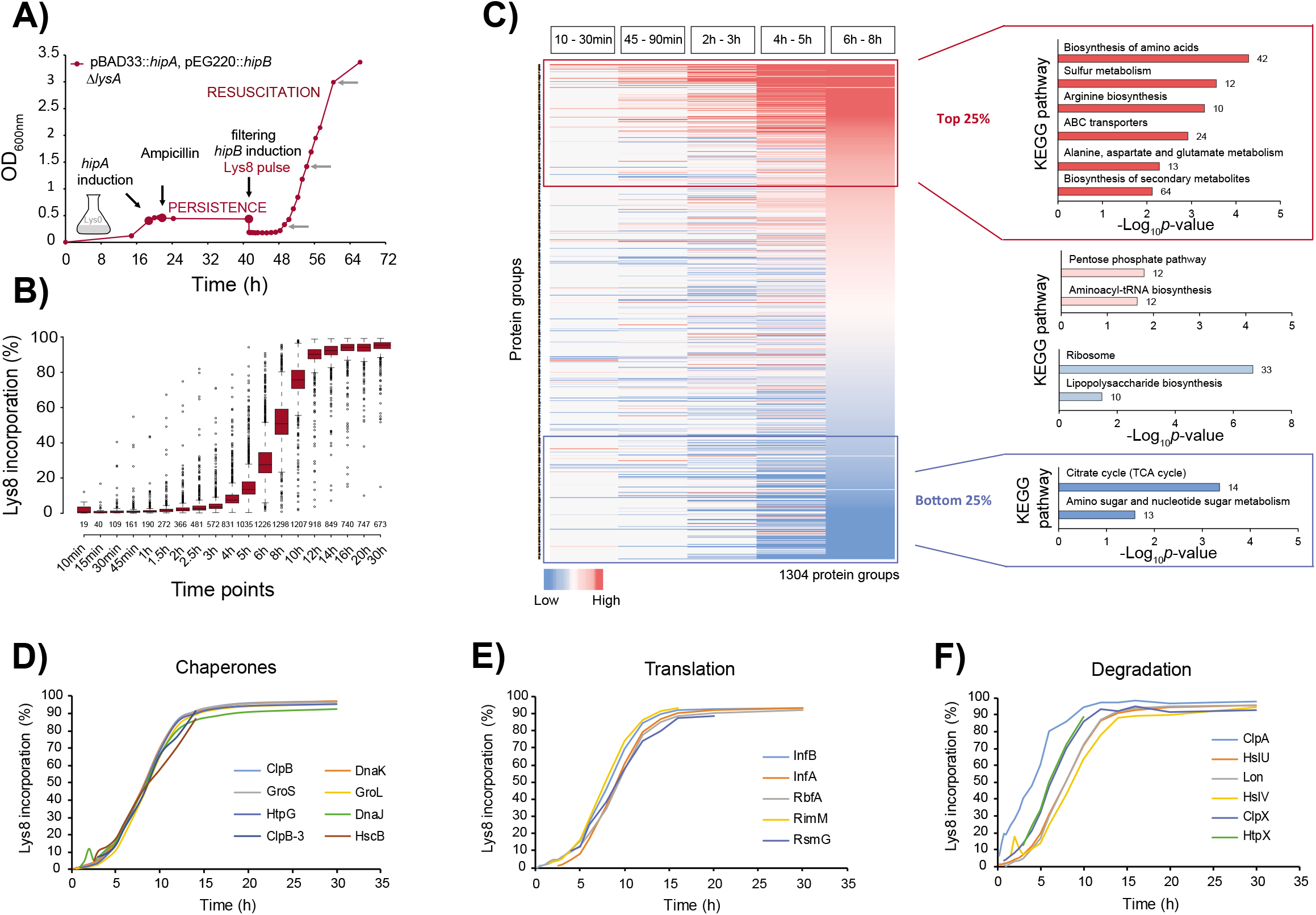
Resuscitating cells restore metabolic activity and translation machinery. **(A)** Transcription of *hipA* was induced and repressed as described in the Legend to Figure 1. To trigger resuscitation, a culture sample was filtered and cells transferred into fresh medium containing IPTG to induce expression of *hipB* (*p_lac_::hipB*) and “heavy” lysine (Lys8) for pulse-labeling. Cells were harvested at 21 time points thereafter. The growth curve is representative of three biological replicates. **(B)** Incorporation of Lys8 into proteins across 19 time points calculated from the measured protein H/L ratios. The box plots are representative of 3 biological replicates. **(C)** Heat map of all quantified proteins color-coded based on their H/L ratio ranking within each time bin. Missing values are colored in grey. KEGG enrichment analysis of proteins in the 25th percentile (Top 25%, red), 25th – 50th percentile (pink), 50th – 75th percentile (light red and light blue) and 75th percentile (Bottom 25%, blue) across all time bins was performed with DAVID software against the background of all quantified proteins. The number of enriched proteins in each category is indicated outside the bars. **(D-F)** Label incorporation curves of selected examples of enzymes involved in protein and folding (chaperones), translation and degradation.

To analyze processes occurring early during resuscitation, we considered time points from the first eight hours of *hipB* induction, in which the optical density doubled. Measured protein ratios were collapsed into five time bins and proteins were ranked based on their H/L ratio within each time bin. According to the KEGG pathway enrichment analysis, proteins exhibiting high label incorporation (top 25%) during the entire eight-hour course of resuscitation are involved mainly in the biosynthesis and transport of amino acids, such as arginine and cysteine, as well as in alanine, aspartate and glutamate metabolism (p-value < 0.05). Conversely, proteins with a low label incorporation (bottom 25%) are constituents of the citric acid cycle, amino sugar and nucleotide sugar metabolism (**Figure 3C**). Altogether, this implies that resuscitating bacteria are predominantly synthesizing enyzmes for anabolic pathways to produce building blocks for cell growth. Proteins involved in catabolic pathways that consume metabolites to release energy are synthesized at a slower rate compared to all quantified proteins. This is in agreement with our experimental design, as we were resuscitating cells with glucose as carbon source. Therefore, most energy was derived from glucose catabolism (glycolysis), not from the catabolism of other metabolites.

### Onset of resuscitation is marked by restoration of glycolysis and translation machinery

To identify proteins that are potentially regulating the switch from persistence to resuscitation, the first time bin (10 – 30min) after induction of *hipB* was analyzed in more detail. GO enrichment analysis revealed that 43 out of 153 quantified proteins take part in translation (*p* = 2.75E-27), out of which 42 are ribosomal proteins and one is the stationary-phase-induced ribosome-associated protein Sra (**Supplementary Table S7,8**). Glycolysis pathway, stress response and protein folding were also significantly enriched. This implies that, at the very onset of resuscitation, cells restored almost the entire translation machinery and restarted their metabolism by inducing glycolysis to convert glucose from the fresh medium into energy used for regrowth (**Supplementary Table S8**). Of note, we recently made a similar observation in resuscitation of chlorotic cyanobacteria [36], pointing to the existence of a common program for awakening of dormant-like cells. Two chaperones (DnaK and GroL) and transcription-related proteins (Rho, DeaD and Pnp) that are involved in RNA metabolism and degradation, were newly synthesized during early response, possibly regulating transcription of specific genes. Because these proteins were also detected to be synthesized during persistence, it is possible that the processes common to persistence are still active during the initial phase of the pulse-labeling. The delay in the wake-up from dormancy was observed previously and depends on several factors, such as the composition of the outgrowth medium or the frequency of persister cells [11, 37]. Several proteins showed unusually high label incorporation in the first 30 minutes after induction of resuscitation. As expected, among them was HipB antitoxin that was induced from the plasmid and therefore served as a positive control. Other proteins participate in various processes, such as the ribonucleotide monophosphatase NagD that dephosphorylates a wide range of (deoxy)ribonucleoside phosphates, or the S-formylglutathione hydrolase FrmB, which converts S-formylglutathione into formate and glutathione to detoxify formaldehyde that can otherwise chemically modify DNA and proteins [38]. Among them were also several proteins known to play a role in persistence, such as the ATP-dependent Clp protease subunit ClpA that directs the ClpAP protease to specific substrates for their degradation [39, 40] (**Figure 3F**) and the RNA polymerase sigma factors RpoS and RpoD. RpoS is important for transcriptional reprograming of many genes that are mainly involved in the metabolism and stress response [40], whereas RpoD preferentially induces transcription of genes associated with fast growth, such as ribosomal proteins [41], which is in agreement with our findings (see below). Considering that RpoD was also detected during persistence, it can be assumed that it might have a dual role during both processes. Moreover, several other proteins were rapidly synthesized during early resuscitation, such as the NemA, which reduces N-ethylmaleimide that inhibits growth by modifying cysteine residues of cellular proteins [42]. NemA is also known to degrade toxic compounds to use them as a source of nitrogen [43](**Supplementary Figure 4**).

### Resuscitation is a concerted and tightly regulated biological process

To get a global insight into processes occurring during resuscitation, we performed statistical enrichment analysis of KEGG and GO-annotated functions in each of the five time bins. Functions related to the ribosome and protein biosynthesis were significantly enriched in all time bins, demonstrating that translation is the key process that dominates resuscitation. In addition, the following protein functions were overrepresented: bin 1 (10-30min): glycolysis and chaperone-mediated protein folding; bin 2 (45-90min): glycolysis and aminoacyl-tRNA synthetase; bin 3 (2-3h): biosynthesis of antibiotics and amino acids; bin 4 (4-5h) and bin 5 (6-8h): biosynthesis of amino acids, antibiotics and carbon metabolism (**Supplementary Table S8**). For 1663 proteins we were able to measure label incorporation over all 21 measured time points. Unsupervised clustering of their temporal profiles revealed eight distinct clusters that differed markedly in the rate of label incorporation, estimated by the average time needed to incorporate 50% of the label (**Supplementary Figure 3, Supplementary Table S9**). Among proteins with fast label incorporation, functions such as “ABC transporters” and “arginine biosynthetic process” were overrepresented (“early functions”). They were followed by protein clusters enriched in diverse metabolic pathways and protein biosynthesis (“intermediate functions”). Proteins with the slowest label incorporation were predominantly involved in various catabolic processes; these “late functions” most likely do not play a role in resuscitation, but may be of importance in transition to stationary phase that was also covered in our analysis. Combined, this analysis revealed that resuscitation is a tightly regulated biological process that starts with activation of protein translation and synthesis of transporters needed to import nutrients and building blocks for protein biosynthesis.

### Ribosomal hibernation factor HflX is induced in resuscitating cells

Almost all detected ribosomal proteins showed synchronized label incorporation during resuscitation (**Figure 4A**). Two notable exemptions were the ribosomal proteins L31-type B and L36-2, that showed markedly higher label incorporation compared to other ribosomal proteins. Approximately 10 hours after *hipB* induction the levels of these two proteins surpassed the levels of their paralogs L31 and L36, pointing to a rearrangement of the large ribosome subunit at the entry of the stationary phase (**Supplementary Figure 2D**). This observation, which was recently reported in another study [44], demonstrates the potential of the used approach to detect dynamics of protein complexes in a rapidly changing biological system. Resuscitation was also marked by induction of HflX that correlated with increased synthesis of ribosomal (and other) proteins (**Figure 4B, Supplementary Table S10**). Of note, the synthesis of other hibernation and stress factors such as RaiA and Sra did not diminish during resuscitation; however, as opposed to persisting cells, they were not in excess compared to newly synthesized ribosomal proteins.

**Figure 4.**
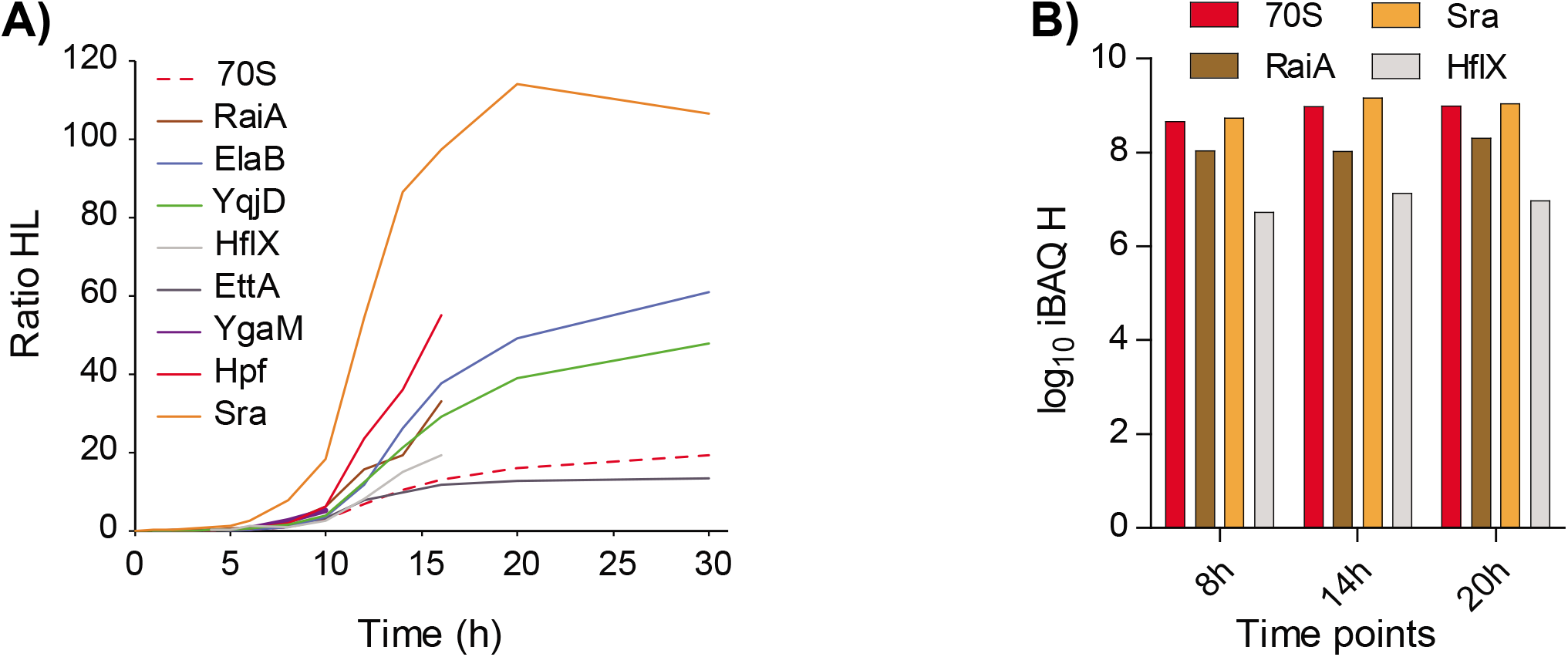
Ribosomal proteins, but also hibernation factors are heavily synthesized in resuscitating cells. **(A)** Label incorporation rates of ribosomal proteins and selected hibernation factors during resuscitation. All ribosomal proteins, except L31-B and L36-2, showed concerted synthesis levels (dotted red trace). The hibernation factor HflX was induced at the onset of resuscitation. Remarkably, ribosome-associated proteins RaiA and Sra were also heavily synthesized. **(B)** Estimates of the amounts of newly synthesized proteins based on iBAQ in the heavy SILAC channel. In contrast to persistence, the amounts of newly synthesized ribosomal proteins are similar to the amounts of newly synthesized RaiA and Sra proteins. The estimated amount of HflX is about 100 times lower.

## Discussion

Numerous studies have shown that bacterial persisters are transcriptionally and metabolically active [8, 12, 45, 46]; however, protein synthesis during antibiotic persistence was so far rarely addressed [15, 27]. Our study complements previous studies by providing a comprehensive and time-resolved analysis of protein synthesis and turnover during antibiotic persistence and, for the first time, during resuscitation. Molecular investigation of persistence is challenging and we note that our approach is not free of experimental bias. Although the model that we used – induction of *hipA* and *hipB* genes of *E. coli* – has been extensively used in previous seminal studies of bacterial persistence [18–23], several uncertainties with this model still remain. Firstly, after *hipA* induction we used the lytic antibiotic ampicillin to enrich for persisters by physically eliminating sensitive cells; this treatment leads to a heterogenous population of cells that includes viable but not culturable (VBNC) cells. Although VBNCs and persister cells are both tolerant to ampicillin and according to some authors present a “dormancy continuum” [47], the exact nature of VBNCs and their relationship to persisting cells is still debated [48]. We note that our present experimental design does not allow us to discriminate between different kinds of persisting cells; however, in future experiments dynamic SILAC can be combined with methods that can separate VBNCs from viable, culturable cells and provide further insights into differences that underline these two cell types. Another, more general question is whether our *hipBA* model in combination with ampicillin treatment is representative of all bacterial persisters. As with every model, this is likely not the case; however, our results should be analyzed and interpreted in the context of numerous previous studies that used the same or similar model to investigate persistence (cited above).

On a technical end, a limitation of our approach is contamination of persister cells with the unlabeled pool of proteins that originate from dead (lysed) cells. Although these proteins can be easily discriminated according to the mass shift of their label, the resulting complexity of the mixture, in combination with a limited dynamic range of mass spectrometric detection, has lead to decreased sensitivity of the measurements and lower proteome coverage compared to a usual shotgun proteomics experiment. This issue could be circumvented by an additional enrichment step to isolate persister cells from the batch culture, for example using long enzymatic treatment that targets the cell membrane [49] or centrifugation [29]. However, such approaches would impose a significant stress to the cells, resulting in proteome changes that could finally lead to data misinterpretation. We cannot exclude that even gentle filtration of the bacterial culture, performed to induce resuscitation and introduce the Lys8 label, introduced some stress to the cell. The alternative could be persister isolation using flow cytometry based on GFP expression from plasmid carrying specific promoters of genes associated with dormancy, as reported previously [23, 50].

Ideally, the enrichment should not be performed on living cells, but rather on isolated proteins that incorporated the label. For this, approaches based on click-chemistry could be used, such as the metabolic labeling with methionine analogue L-azidohomoalanine (AHA) with subsequent attachment of an affinity tag to the reactive azide-group and enrichment of the labeled proteins by affinity chromatography [51]. However, in our experience the application of AHA interferes significantly with translation and causes severe growth defects, which is not compatible with our study.

As mentioned above, application of labeled amino acids to study protein synthesis and turnover is often hampered by a limited dynamic range of detection in a mass spectrometer. This means that very low level of label incorporation will not be detected in a high background of MS signals originating from unlabeled peptides, making detection of label incorporation more likely in abundant proteins. Although the reasons for this are not only technical, as proteins involved in translation and essential proteins are usually abundant, this could be ameliorated by addition of stable isotope-labeled peptide standards or even the whole cell lysates like reported in previous analyses of low abundant post-translational modifications [52]. A further complication concerns the recycling of amino acids released from degradation of pre-existing proteins into the medium, which causes a dilution of labeled amino acid pool with unlabeled amino acids and results in apparently lower turnover rates that cannot truly describe actual values. These values can be corrected using a recycling factor that can be calculated by measuring label incorporation in partially labeled missed cleaved peptides [24]. Although these issues may influence the overall sensitivity and accurate estimation of individual protein turnover rates, we do not expect that they lead to a false detection of the newly synthesized proteins.

## Materials and Methods

### Bacterial strains and plasmids

*E. coli* strains and plasmids used in this study are listed in the **Table 1**. Due to a high toxicity of HipA protein, *hipA* gene was cloned into the expression plasmids together with a Shine-Dalgarno sequence in which the spacer between the Shine-Dalgarno and the start codon was changed to decrease the translation efficiency of HipA [53]. In Table S1, sd8 specifies a consensus sequence AAGGAA with a spacer of eight nucleotides to the GTG start codon. Oligonucleotides used are listed in the **Table 2**.

**Table 1.** List of bacterial strains and plasmids.

| Strains<br>Plasmids | Genotype | Source |
| --- | --- | --- |
| **Strains** |  |  |
| **MG1655** | Wild-type *E. coli* | [56] |
| ***ΔlysA*** | MG1655 *ΔlysA::FRT* P1 transduction from BW25113<br>*lysA::FRT::aphA::FRT* (KEIO collection) into MG1655 and flipped out. | This work |
| ***sra::kana<sup>R</sup>*** | MG1655 *sra::FRT::aphA::FRT* P1 transduction from BW25113<br>*sra::FRT::aphA::FRT* (KEIO collection) into MG1655 | This work |
| ***raiA::kana<sup>R</sup>*** | MG1655 *raiA::FRT::aphA::FRT* P1 transduction from BW25113<br>*raiA::FRT::aphA::FRT* (KEIO collection) into MG1655 | This work |
| **Plasmids** |  |  |
| **pBAD33** | p15 derived expression plasmid, *cat*, *araC*, P<sub>BAD</sub> promoter, Cm<sup>R</sup> | [57] |
| **pNDM220** | R1 derived expression plasmid, p<sub>A1/O4/O3</sub> (p<sub>lac</sub>), *lacI<sup>q</sup>*, *aphA* (Km<sup>R</sup>) | [55] |
| **pGOOD** | pTrcHisB and pACYC184 derived plasmid, *lacI*, TetR | [58] |
| **pBAD33::hipA (pEG5)** | pBAD33 P<sub>BAD</sub>:: *sd8 gtg hipA* | [18] |
| **pEG220::hipB** | pEG220 P<sub>lac</sub>:: *sdopt::hipB*, Km<sup>R</sup> | This work |
| **pGOOD::psra** | pGOOD, *sra* and its native promoter cloned under *lacI* promoter upon digestion with BglII | This work |

The *lysA::FRT::Kan::FRT* deletion strain from the Keio collection [54] was P1 transduced into MG1655. The resulting mutation was verified by PCR diagnostics using primers EG239 and EG240 and then transformed with pCP20 to induce the flippase to flip out the *Kan^R^* gene, resulting in MG1655 *lysA::FRT* mutant. The *sra::FRT::Kan::FRT* and The *sra::FRT::Kan::FRT* deletion strains from the Keio collection [54] was P1 transduced into MG1655.

**Table 2.**
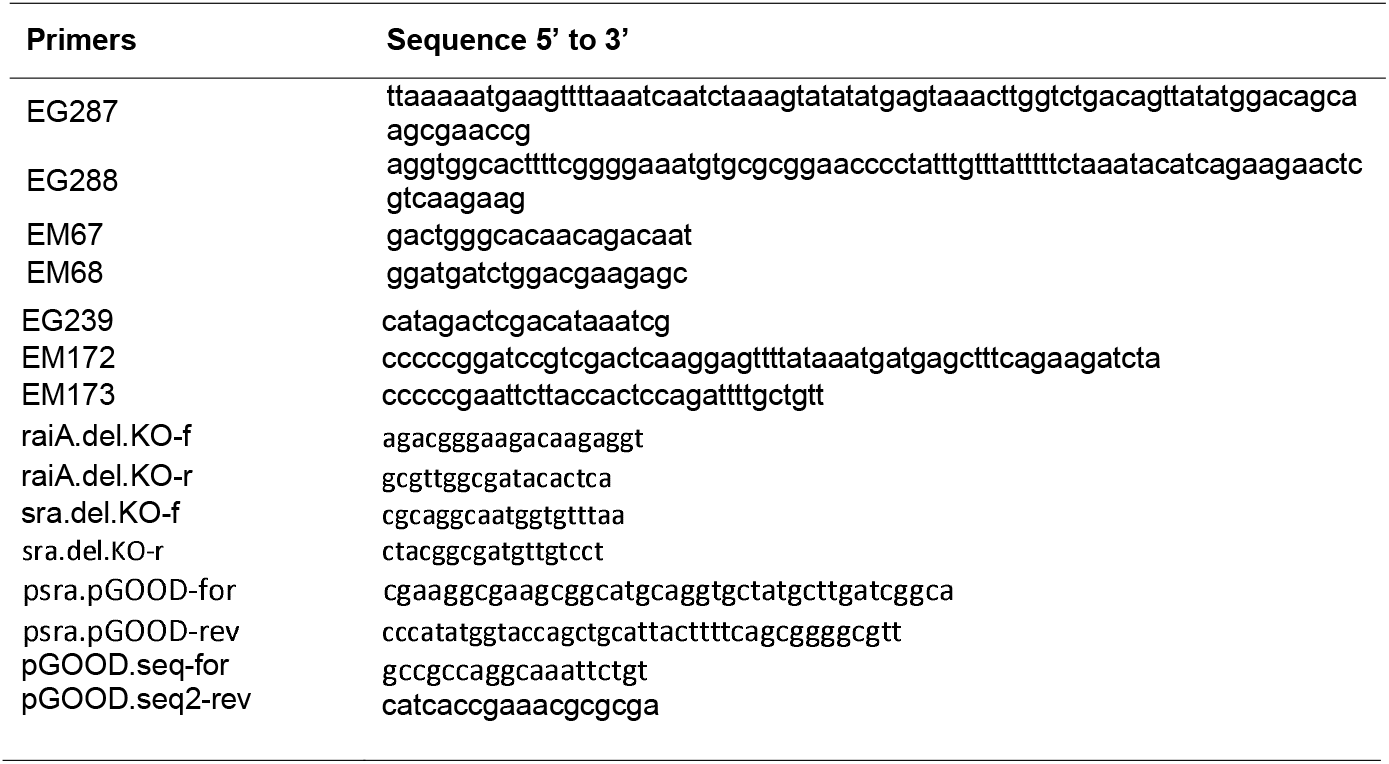
List of DNA oligonucleotides.

Plasmid pEG220 was constructed to change the kanamycin cassette of the expression plasmid pNDM220 [55] to ampicillin. The ampicillin cassette has been amplified with primer EG287 and EG288 from pKD4. Then, DY331 strain has been electroporated with PCR fragment following the procedure from D.Court’s laboratory (https://redrecombineering.ncifcrf.gov/protocols/). The correct insertion of the ampicillin cassette has been verified using primer EM67 and EM68 matching in the insertion in the pNDM220. Then, the kanamycin cassette and the surrounding region of the newly constructed pEG220 were sequenced using primers EM67 and EM68.

For construction of plasmid pEG220::hipB, *hipB* coding region with an optimized Shine Dalgarno was PCR amplified using primer EM172 and EM173, the resulting DNA fragment digested with BamHI and EcoRI and ligated in pEG220 digested with the same enzymes.

For construction of plasmid *pGOOD::psra*, plasmid was digested with BGlI and Mph1103I, fragment conteining original lac promoter was removed and fragment containing *sra* gene and its native promoter, amplified with psra.pGOOD-for and rev, was cloned. Plasmid was sequenced wit pGOOD.seq-forand pGOODseq2-rev.

### Persistence analysis

Pre-cultures were grown overnight in liquid medium (Luria-Broth or M9 minimal medium) supplemented with 0.4% (w/v) glucose, 25 μg/ml chloramphenicol for maintenance of pBAD33::*hipA* plasmid, for *sra::kana^r^* and *raiA::kana^r^*, additional 50 μg/ml kanamycin. Day cultures were inoculated at OD_600nm_ of 0.08; at OD_600nm_ of 0.3 HipA was induced with 0.2% (w/v) L-(+)-arabinose for 1h, followed by 100 μg/ml ampicillin (Figure 2). For analysis in M9, bacteria grew till at OD_600nm_ of 0.4 and induced for 3h with 0.2 % (w/v) L-(+)-arabinose, followed by 100 μg/ml ampicillin (Supplementary Figure 1). To analyse persistor under native HipA expression level (Figure 2), day culture were inoculated in LB at OD_600nm_ of 0.08 followed by 100 μg/ml ampicillin addition at OD_600nm_ of 0.5. Cells were harvested at desired time points (1ml), pellets were washed with phosphate-saline buffer (PBS), serially diluted and spotted on LB-agar plates supplemented with 0.4% (w/v) glucose. Experiments were performed in triplicates. Data were visualized using Prism 8 (GraphPad).

### P1 phage preparation and transduction

Pre-culture were inculated in LB broth supplemented with 50 μg/ml of kanamycin. For lysate preparation, the overnight donor culture was diluted 1:100 into 5 ml of 0.2% (w/v) glucose and 5 mM CaCl2 and incubated 30 to 45 min at 37°C with shaking. 100 μl of P1 phage stock were added to the growing culture and incubated, with shaking, until the culture lysed (~3h). To complete cell lysis few drops of CHCl_3_ were added to the shaking culture. The culture was pelleted via centrifugation for 10 min at ~9200 × g at 4°C and the supernatant containing the phages was filtered over a 0.45 μm sterile filter. For P1 transduction, *E. coli* recipient strain was inoculated in 5ml LB broth overnight at 37° C. The following day, 1.5 ml of the cell culture was pelleted by centrifugation for two minutes at maximum speed at room temperature and the pellet was resuspended in P1 salt solution (10 mM CaCl_2_, 5 mM MgSO_4_), at half of the original volume. 100 μl of the cell/P1 mixture was mixed with different amounts of the desired P1 lysate (1, 10, and 100 μl) and the phage was allowed to adsorb for 30 min at room temperature. One mililiter LB broth plus 200 μl of 1 M sodium citrate was added to the cell/phage mix and incubated for 1h at 37° C. Cultures were then centrifuged, supernatants removed and the pellets resuspended in LB medium. Resuspended cells were plated on LB agar plates supplemented with 50 μg/ml kanamycin. Single colonies were streaked on fresh selective plates and confirmed by PCR.

### Cell growth and SILAC labeling

Cells were grown in 100 ml of M9 minimal medium (50 mM Na_2_HPO_4_, 22 mM KH_2_PO_4_, 8.6 mM NaCl, 18.7 mM NH_4_Cl, 1 mM MgSO_4_, 0.1 mM CaCal_2_, 0.0001% (w/v) thiamine) supplemented with 0.4% (v/v) glycerol, 25 μg/ml chloramphenicol for maintenance of pBAD33::*hipA* plasmid and 50 μg/ml kanamycin for maintenance of pEG220::*hipB* plasmid. Cultures were grown in batch at 37°C shaking at 200 rpm. Pre-cultures were grown for 20 hours in M9 medium supplemented with 0.4% (w/v) D-(+)-glucose to repress the P_BAD_ promoter. Main cultures were grown starting from OD_600nm_ of 0.01 – 0.02 in M9 medium without glucose. For HipA-induced persistence, wild type *E. coli* K-12 MG1655 strain was used. The expression of *hipA* was induced with 0.2% (w/v) L-(+)-arabinose at OD_600nm_ of ≈0.4. After 3 hours of *hipA* expression, cultures were treated with 100 μg/ml ampicillin for 19 - 22 hours to kill non-persister cells and 0.4% glucose to repress transcription of the P_BAD_::*hipA* promoter fusion. A pulse of 0.025% (w/v) stable isotope-labeled lysine derivative ^13^C_6_^15^N_2_ L-lysine (“heavy” lysine, Lys8, K8, Cambridge Isotope Laboratories) was then added in the presence of ampicillin and 1 – 2 mL of cultures were harvested in intermittent time intervals (10min - 24h) by centrifugation at 4°C. A time point before the pulse of Lys8 (0h) was harvested as a control. For the resuscitation experiment, an MG1655 derived strain lacking *lysA (ΔlysA)* was used. Both pre-cultures and main cultures were supplemented with 0.025% (w/v) natural L-lysine (“light” lysine, Lys0, K0, Sigma Aldrich). The expression of *hipA* was induced with 0.2% (w/v) arabinose at OD_600nm_ of ≈0.4. After 3 hours of *hipA* expression, cultures were treated with 0.4% glucose and 100 μg/ml ampicillin for around 22 hours. To induce resuscitation, cultures were first quickly filtered using pre-warmed 1 l Corning filter system (0.22 μm pore size) to remove ampicillin, Lys0 and dead cells. Immediately after, cells were quickly recovered in pre-warmed medium supplemented with 25 μg/ml chloramphenicol, 50 μg/ml kanamycin, 0.4% glucose, 0.025% Lys8 and 2 mM β-D-1-thiogalactopyranoside (IPTG) for the induction of the *P_lac_::hipB* fusion. Cells were harvested at intermittent time intervals (10min - 30h) by centrifugation at 4°C. A time point before filtering and Lys8 pulse (0h) was harvested as a control.

### Cell lysis and sample preparation

Collected cell pellets were lyzed in a lysis buffer (40 mg/ml SDS, 100 mM Tris-HCl pH 8.6, 10 mM EDTA and Complete protease inhibitors (Roche)) and sonicated 3 times for 30 s at 40% amplitude. The cellular debris was pelleted by centrifugation at 15,000 g for 30 min at room temperature and proteins were precipitated from the supernatant with chloroform and methanol. Protein pellets were dissolved in a denaturation buffer (6 M urea, 2 M thiourea and 10 mM Tris pH 8.0). Protein concentration was determined using standard Bradford assay (Bio-Rad). A 10 μg protein aliquot from each time point was reduced using 1 mM dithiotreitol for one hour and alkylated with 5.5 mM iodoacetamide for additional hour. Proteins were pre-digested with endoproteinase Lys-C (1:100 w/w) for 3 hours, then diluted with 4 volumes of 20 mM ammonium bicarbonate, pH 8.0 and digested overnight with Lys-C (1:100 w/w) at room temperature. The reaction was stopped by lowering pH to 2 with trifluoroacetic acid. Peptides were purified via stage tips [59] as described previously [19]. Briefly, before each LC-MS/MS measurement, samples were de-salted and purified on reversed-phase C18 stage tips (Empore), inhouse prepared, previously activated with methanol and equilibrated with solvent A*. Up to 10 μg of peptides were loaded onto the membrane, washed with solvent A and then eluted with 50 μl of solvent B [80% (v/v) acetonitrile and 0.1% (v/v) formic acid] and concentrated by vacuum centrifugation. The sample volume was adjusted with solvent A and final 10% (v/v) of solvent A*.

### LC-MS/MS measurement

Samples from each time point were separated by the EASY-nLC 1200 system (Thermo Scientific). Peptide separation was performed by reversed-phase chromatography on the in-house packed analytical column (20 cm × 75 μm, 1.9 μm ReproSil-Pur C18-AQ particles (Dr. Maisch)). Peptides were loaded onto the column at a flow rate of 700 nl/min of solvent A (0.1% (v/v) formic acid) and a constant temperature of 40°C under maximum back-pressure of 850 bar. Peptides were eluted using 116 min segmented gradient of 10 – 50% solvent B (80% (v/v) acetonitrile, 0.1% (v/v) formic acid) at a flow rate of 200 nl/min. Peptides were infused directly from the column tip into the on-line coupled Q Exactive HF mass spectrometer (Thermo Scientific) using nanoelectrospray ion source (Thermo Scientific) at a voltage of 2.3 kV and a temperature of 275 °C. Positively charged peptides were analyzed in a data-dependent acquisition mode, in which one full scan and subsequent MS/MS scans of 12 (Top12 method) most abundant precursors (isolation window of 1.4 m/z) were recorded in a mass range from 300 to 1,650 m/z. To prevent repeated fragmentation, the masses of sequenced precursors were dynamically excluded for 30 s. Only precursors with assigned charge states ≥ 2 and ≤ 5 were considered for fragmentation selection. Full scans were acquired with a resolution of 60,000 at m/z 200, the maximum injection time of 25 ms and the automatic gain control (AGC) target value of 3 × 10^6^ charges. The higher energy collisional dissociation (HCD) fragmentation was achieved at normalized collision energy of 27% and intensity threshold of 1 × 10^5^. MS/MS spectra were acquired with the resolution of 30,000, the maximum injection time of 45 ms and the AGC target value of 1 × 10^5^ charges.

### MS data processing and analysis

Acquired raw data was processed with MaxQuant software (version 1.5.2.8) [60] using the default settings if not stated otherwise. Raw files of particular experiments were processed separately. The built-in Andromeda search engines searched MS/MS spectra against fragment masses of peptides derived from an *E. coli* K12 reference proteome (taxonomy ID 83333) containing 4,324 entries (UniProt, release 2017/12) and a list of 245 common contaminants. The minimum required peptide length was set to 7 amino acids with the maximum of two missed cleavage sites allowed for endoproteinase Lys-C. For a protein to be quantified, two occurrences of the protein H/L ratio were required per time point (the multiplicity was set to two with Lys8 specified as the heavy label). Carbamidomethylation of cysteine was set as a fixed modification, and protein N-terminal acetylation and methionine oxidation as variable modifications. Precursor ion mass tolerance was set to 20 ppm in the first search and to 4.5 ppm and in the main search. Peptide and protein identifications were filtered using a target-decoy approach with a an estimated false discovery rate (FDR) of less than 1.3% at protein and less than 0.4% at peptide level[61]. Proteins identified by the same set of peptides were combined into a single protein group. For protein quantification, at least 2 peptide ratio counts were required.

Data analysis was performed using Microsoft Excel and Perseus software (version 1.5.6.0)[62]. Nonnormalized protein H/L ratios from proteingroup.txt were used for protein quantification. All contaminants, reversed hits and proteins identified only by modification were removed. To simplify the data, the data sets from three biological replicate measurements were combined as a union, in which protein H/L ratios in each of the time points were kept if measured only in one replicate (Class I), a mean was calculated if measured in two (Class II) or in three replicates (Class III). To reduce the number of data points and missing values, time points were pooled into time bins. For that, the median protein H/L ratio of respective time points was calculated and assigned to a specific time bin. Proteins were then ranked based on their H/L ratio within each time bin and displayed in a heatmap-like representation sorted by the maximal H/L ratio across time bins. Gene-annotation and KEGG enrichment analysis was performed using DAVID tool (version 6.8) with default parameters and the background of all quantified proteins in particular experiment [63].

### Temporal analysis of label incorportion

MaxQuant processed data were imported in the R environment (R Core Team, 2018) to perform a time series analysis. The heavy label incorporation was calculated by dividing the heavy label intensity by the total (heavy plus light) intensity for each protein. Reverse hits, potential contaminants and proteins only identified by sites were filtered out. The median of three replicates was computed per time points. Proteins with missing values were discarded from further analysis. The Euclidean distance between proteins was computed and used to generate the hierarchical clustering via the Ward’s minimum distance using the stats R package. Time period resolution was achieved with minimal overlap when using eight cluster groups (k = 8). The resulting heatmap was drawn using the gplots R package (Warnes et al., 2019), while cluster protein profiles over time were generated using the ggplot2 R package (Wickham, 2016). For each cluster, the time needed to reach 50% heavy label incorporation was inferred by intersecting the 50% incorporation with the smoothed conditional mean. The list of proteins belonging to each cluster were imported into Perseus software suite (version 1.6.5.0) (Tyanova et al., 2016). The proteins were functionally annotated with Gene Ontology, KEGG, Pfam, GSEA, UniProt keywords, Corum, Interpro, Reactome and EC functional resources. Each cluster was tested for function categories over- or under-representation using a fisher exact test (FDR ≤ 0.1).

### Reactive web application for data browsing

To allow the scientific community to browse our datasets, we developed a Shiny application coded entirely in the R programming language (Chang et al., 2019). Our application, called PCTsee, allows user to load different datasets, which are available through a drop down widget. It is then possible to select gene name or protein ID of interest in order to display their abundance value. Whereby the abundance values have been computed by the MaxQuant software during LC-MS/MS data processing (e.g. intensity, ratio). The user selection will result in a protein abundance profile plot, as well as a table containing some information about the selected protein (e.g. protein names, sequence coverage). Users can interact further with the displayed plot to zoom, rescale or remove samples due to the plotly R package (Sievert, 2018). PCTsee application is available at the following address: https://pctsee.pct.uni-tuebingen.de.

### GO_term enrichment

Gene ontology analysis of the proteins actively synthesized during resuscitation was perform with the online software DAVID (https://david.ncifcrf.gov/tools.isp). Enrichment was performed comparing the gene names quantified proteins against the ones of identified proteins. DAVID was used with its preset with: similarity overlap 3 and threshold 0.5; initial and final group membership 3 and multiple linkage 0.5; enrichment thresohold at EASE 1.

### Calculation of protein turnover rates

Protein turnover rate (*k*) was calculated by linear regression of natural logarithm of protein SILAC H/L ratio over time as described previously [24] using equation independent of the growth rate:

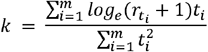

Where *m* is the number of time points (*t_i_*) and *r_t_i__*. is protein H/L ratio measured in a time point *t_i_*. The half-life of a protein (*T*_1/2_) was calculated by dividing the natural logarithm with the turnover rate *k*:

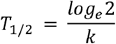

Protein H/L ratios derived from a union of three replicates were used for the calculation and only proteins with H/L ratio measured in > 4 time points were considered. As a quality check of linear regression, the coefficient of determination (R^2^) was required to be > 0.70 to ensure good curve fitting and reliable turnover rate estimation. All proteins with determined protein turnover rates are reported in the **Supplementary Table S5**.

The mass spectrometry proteomics data have been deposited to the ProteomeXchange Consortium via the PRIDE [64] partner repository with the dataset identifier PXD018153. All other data needed to evaluate the conclusions in this paper are present in the paper or the Supplementary Materials.

## Supplementary Material

**Supplementary Figure 1.**
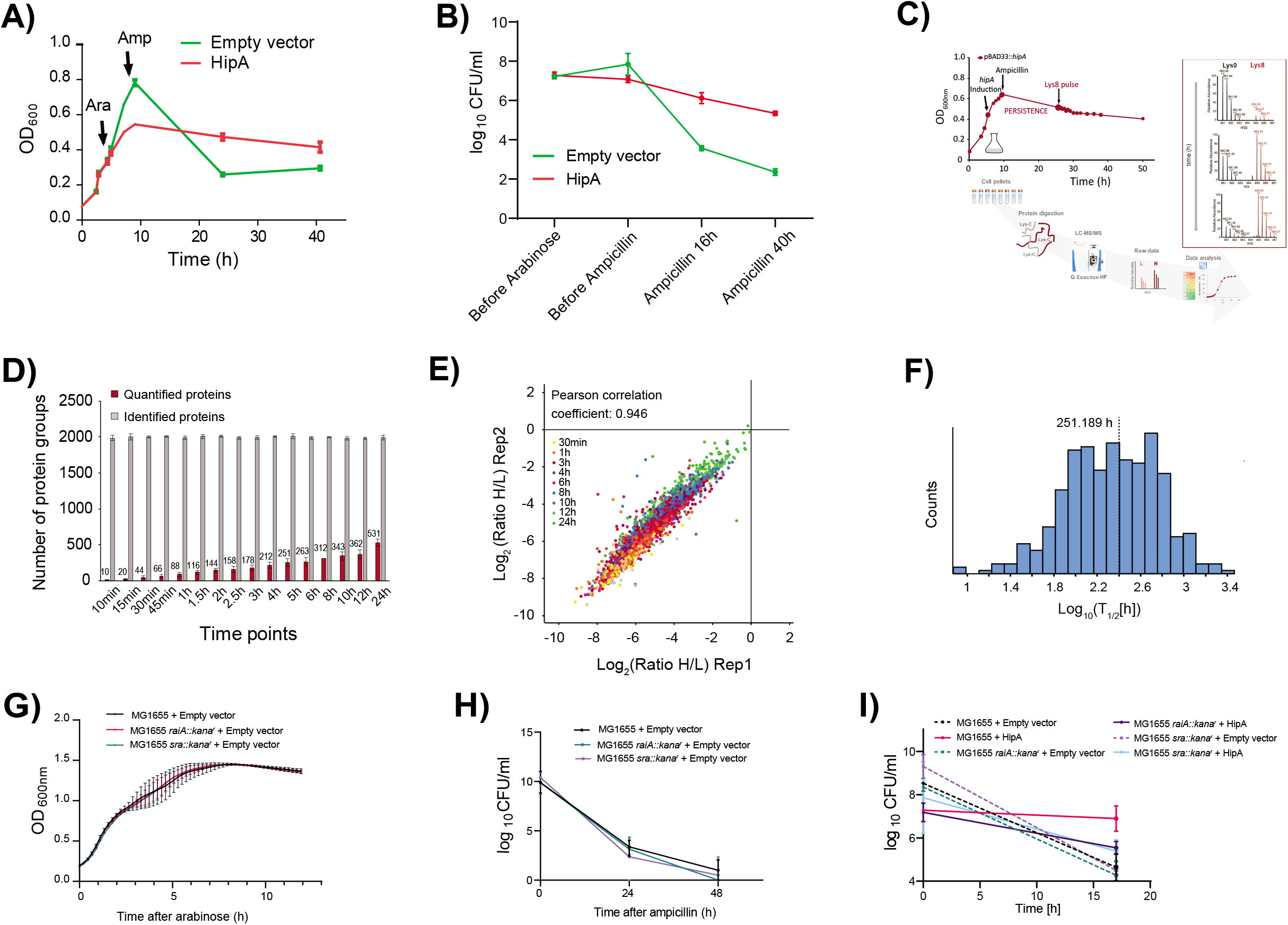
Measurement of newly synthesized proteins in persisting *E. coli* cells. **(A)** OD_600_ measurement of the *E. coli* culture after induction of ectopic hipA expression compared to the empty vector; **(B)** Colony-forming units (CFU) measurement of the hipA-expressing *E. coli* culture at several points before and after ampicillin addition; *E. coli* cells containing empty vector were used as control; **(C)** Biochemical and mass spectrometry workflow employed in our study; **(D)** Lys8 label incorporation monitored by increasing number of quantified proteins over 24 hours. Quantified proteins were detected in both “light” (“L”) and “heavy” (“H”) form, which enabled calculation of their H/L ratios (relative quantification); **(E)** Correlation plot demonstrates high reproducibility between two biological replicates; **(F)** Estimated half-lives of newly synthesized proteins during persistence; **(G)** Empty vector controls for HipA growth effect measurements in LB medium (**Figure 2C**); **(H)** Empty vector controls for persistence measurements in LB medium (**Figure 2D**); **(I)** CFU measurement of *E. coli* wild type and strains lacking of *sra* and *raiA* before and after ampicillin addition, in M9 medium.

**Supplementary Figure 2.**
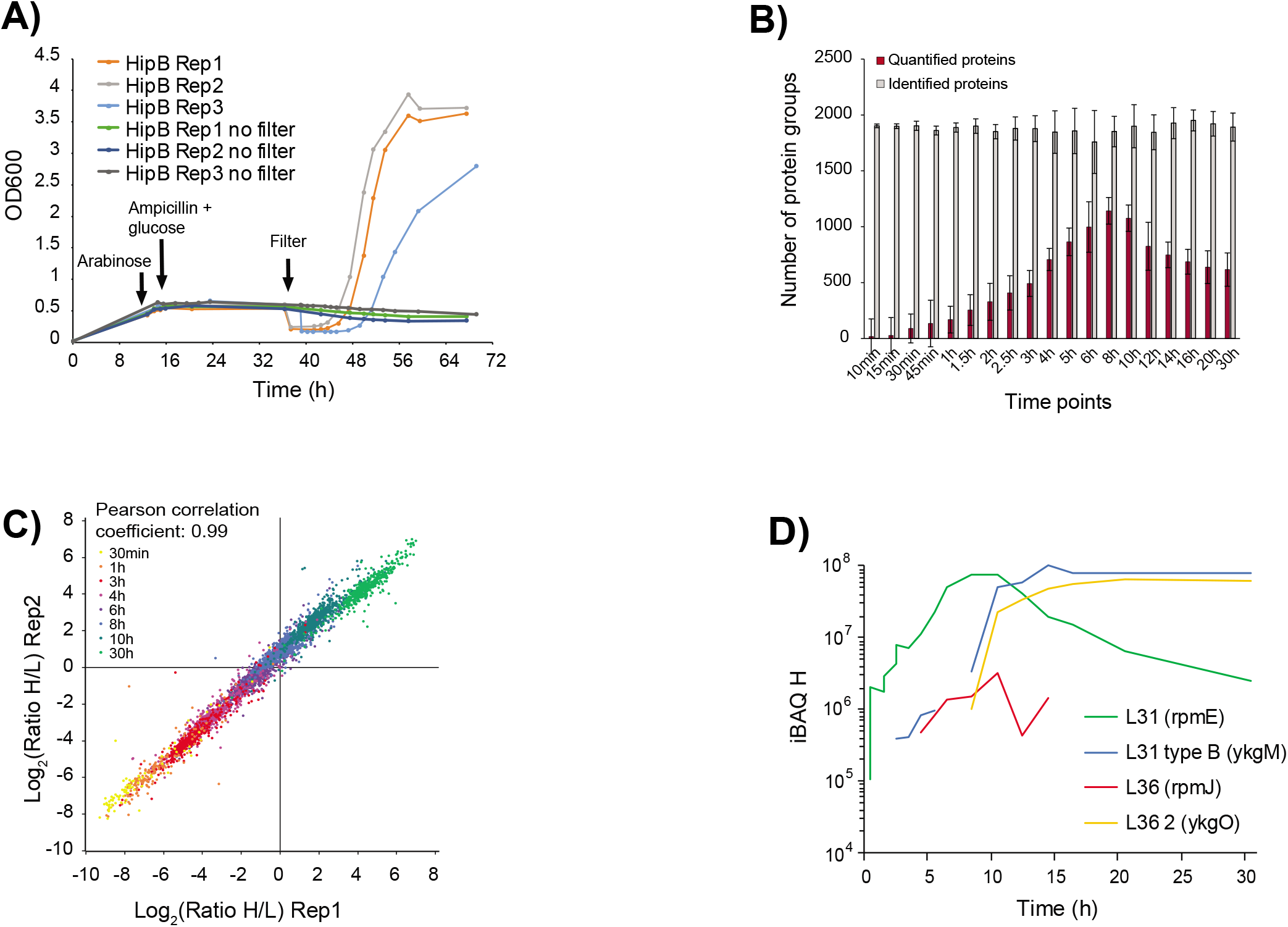
Measurement of newly synthesized proteins in resuscitating *E. coli* cells. **(A)** OD_600_ measurements of *hipB*-induced and -uninduced (“no filter”) cultures confirm that resuscitation was driven by ectopic *hipB* expression. In the absence of hipB expression the OD_600_ remained constant for at least 30 hours, demonstrating that no spontaneous resuscitation or development of resistance occurred during this time period. (B) Lys8 label incorporation monitored by increasing number of quantified proteins over 30 hours. Note that after eight hours cell entered the exponential growth, which led to exponential increase of the heavy signal intensity. After this time point numerous proteins are predominantly present in their heavy form and their H/L ratio cannot be calculated, which leads to an apparent “decrease” of the quantified proteins. **(C)** Correlation plot of two biological replicates demonstrates high reproducibility. **(D)** Abundance traces of ribosomal proteins L31 (*rpmE*) and L36 (*rpmJ*), and their paralogs L31 type B (*ykgM*) and L36 2 (*ykgO*) point to a rearrangement of the large ribosomal unit at the entrance to stationary phase.

**Supplementary Figure 3.**
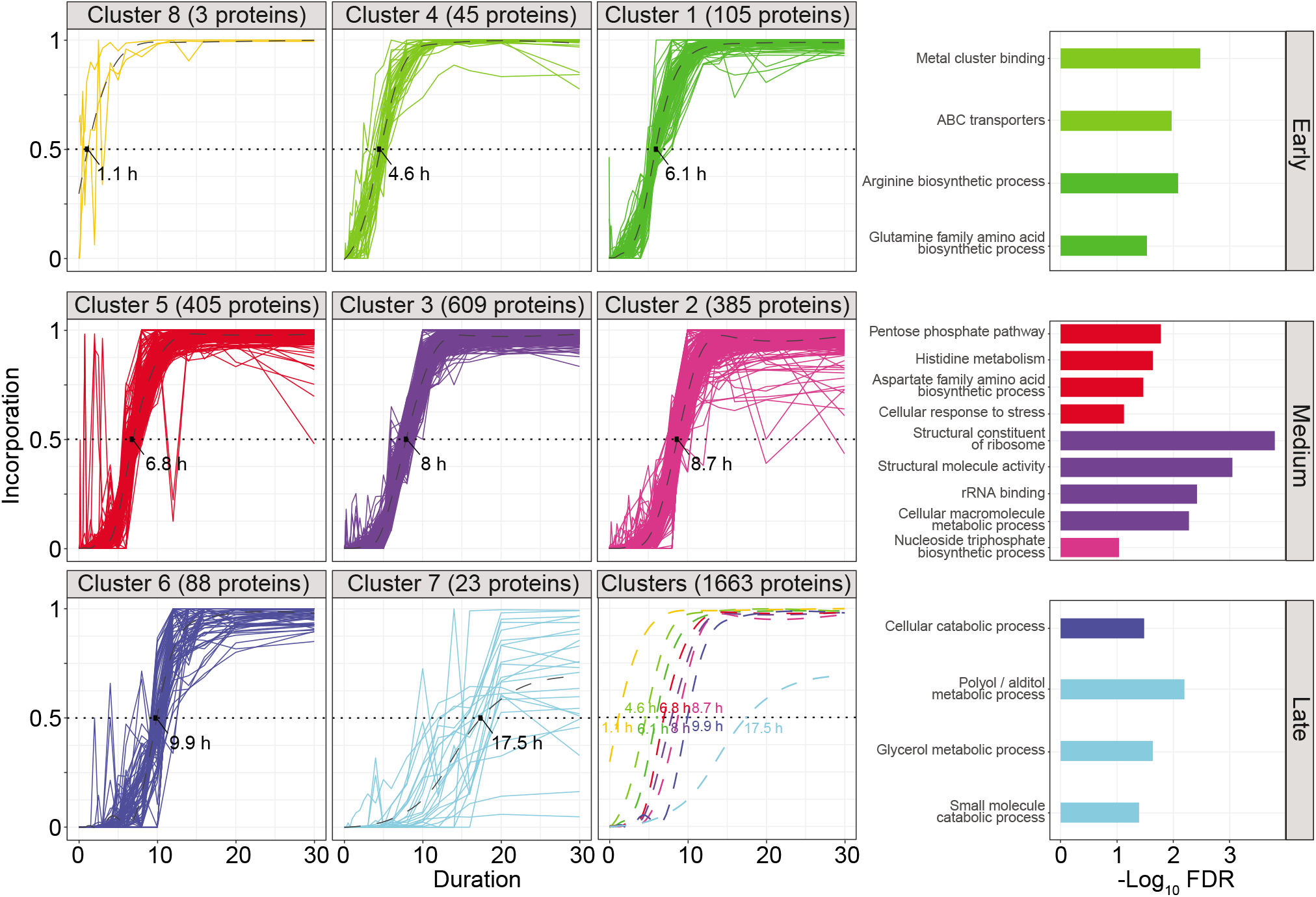
Clustering of label incorporation profiles allows reconstruction of time-dependant cell metabolism during resuscitation. Heavy label incorporation was used to cluster proteins across the resuscitation time scale. Among the eight clusters represented, clusters 8, 4 and 1 belong to the early synthesized proteins; clusters 5, 3 and 2 belong to the medium proteins; and clusters 6 and 7 belong to the late proteins. The smoothed conditional means is represented as a dashed curve within each cluster. For each cluster the time at 50% incorporation is determined based on the intersection with smoothed means. The top significantly over-represented functional category for each cluster (FDR ≤ 0.05) are plotted on the right panels.

**Supplementary Figure 4.**
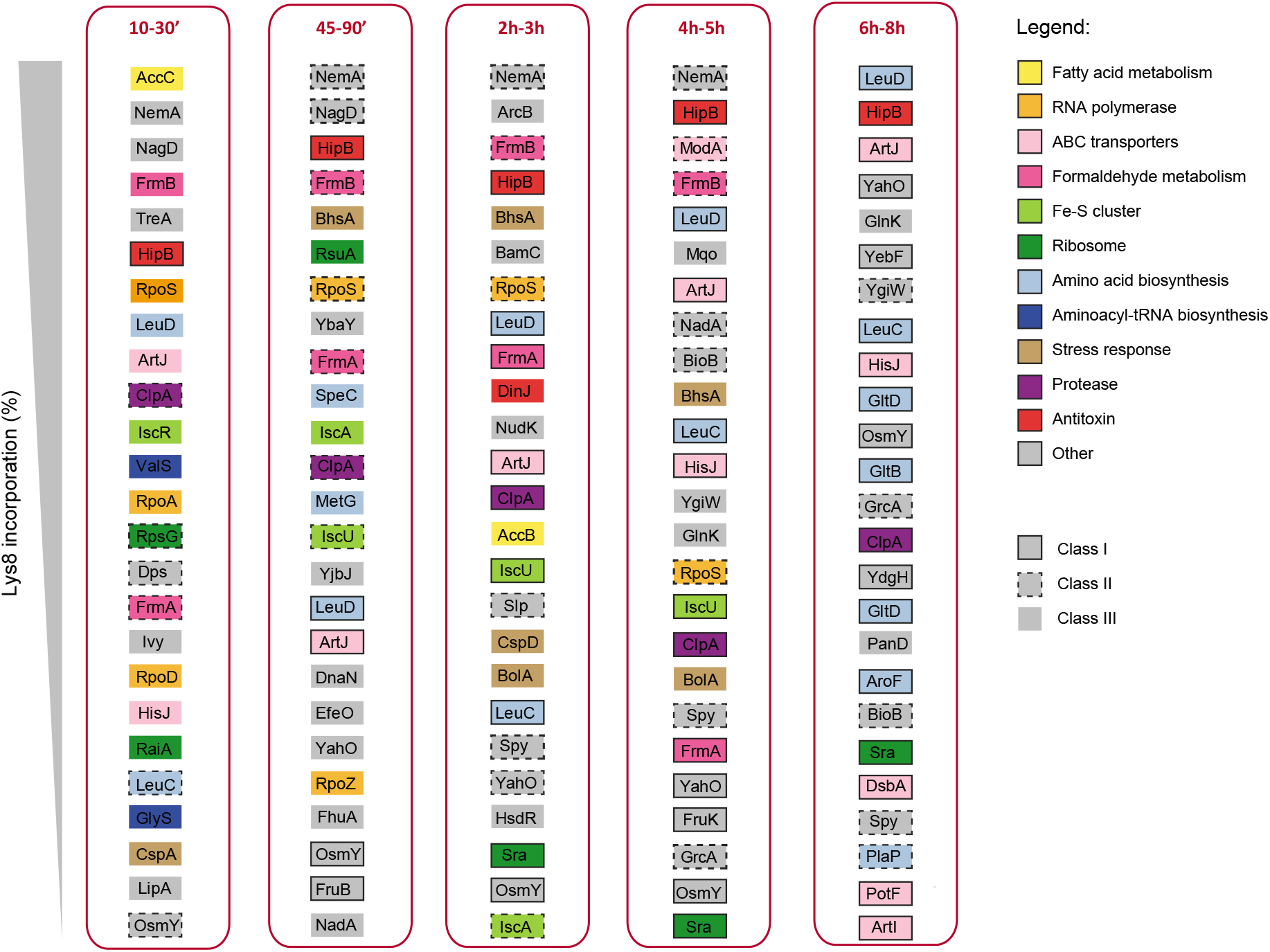
Map of top 25 quantified proteins with highest incorporation of Lys8 in each time bin during resuscitation. Different functional descriptions are color coded according to the legend. Boxes around individual proteins are coded based on the style of the line and indicate the confidence of protein quantification per time bin according to the following classification system: protein is quantified in > 67% of the time points in 3 replicates (Class I), protein is quantified in < 67% and > 33% of the time points in 3 replicates (Class II), protein is quantified in < 33% of the time points in 3 replicates (Class III).

## Acknowledgements

The authors thank Karl Forchhammer for useful discussions and comments on the manuscript. B.M. was supported by grants from the Deutsche Forschungsgemeinschaft (German Research Foundation Cluster of Excellence EXC 2124, SFB 766, FOR 2816, TRR 261), the European Union’s Horizon 2020 research and innovation programme under the Marie Sklodowska-Curie grant agreement No. 955626 and the German-Israeli Foundation grant I-1464-416.13/2018. K.G. was supported by a grant from the Danish National Research Foundation (grant identifier DNRF120) and the Novo Nordisk Foundation.

## References

1. Ventola, C.L., The Antibiotic Resistance Crisis: Part 1: Causes and Threats. Pharmacy and Therapeutics, 2015. 40(4): p. 277–283.

2. Brauner, A., et al., Distinguishing between resistance, tolerance and persistence to antibiotic treatment. Nature Reviews Microbiology, 2016. 14:p. 320.

3. Van den Bergh, B., M. Fauvart, and J. Michiels, Formation, physiology, ecology, evolution and clinical importance of bacterial persisters. FEMS Microbiology Reviews, 2017. 41(3): p. 219–251.

4. Balaban, N.Q., et al., Bacterial Persistence as a Phenotypic Switch. Science, 2004. 305(5690): p. 1622–1625.

5. Lewis, K., Persister Cells. Annual Review of Microbiology, 2010. 64(1): p. 357–372.

6. Balaban, N.Q., Persistence: mechanisms for triggering and enhancing phenotypic variability. Current Opinion in Genetics & Development, 2011. 21(6): p. 768–775.

7. Balaban, N.Q., et al., Definitions and guidelines for research on antibiotic persistence. Nat Rev Microbiol, 2019. 17(7):p. 441–448.

8. Amato, S.M., et al., The role of metabolism in bacterial persistence. Frontiers in Microbiology, 2014. 5: p. 70.

9. Page, R. and W. Peti, Toxin-antitoxin systems in bacterial growth arrest and persistence. Nature Chemical Biology, 2016. 12:p. 208.

10. Harms, A., E. Maisonneuve, and K. Gerdes, Mechanisms of bacterial persistence during stress and antibiotic exposure. Science, 2016. 354(6318).

11. Jõers, A., N. Kaldalu, and T. Tenson, The Frequency of Persisters in Escherichia coli Reflects the Kinetics of Awakening from Dormancy. Journal of Bacteriology, 2010.192(13): p. 3379–3384.

12. Orman, M.A. and M.P. Brynildsen, Establishment of a method to rapidly assay bacterial persister metabolism. Antimicrob Agents Chemother, 2013. 57(9): p. 4398–409.

13. Orman, M.A. and M.P. Brynildsen, Dormancy is not necessary or sufficient for bacterial persistence. Antimicrob Agents Chemother, 2013. 57(7): p. 3230–9.

14. Roostalu, J., et al., Cell division in Escherichia coli cultures monitored at single cell resolution. BMC Microbiol, 2008. 8: p. 68.

15. Radzikowski, J.L., H. Schramke, and M. Heinemann, Bacterial persistence from a system-level perspective. Curr Opin Biotechnol, 2017. 46:p. 98–105.

16. Rowe, S.E., et al., Persisters: Methods for Isolation and Identifying Contributing Factors--A Review. Methods Mol Biol, 2016. 1333:p. 17–28.

17. Canas-Duarte, S.J., S. Restrepo, and J.M. Pedraza, Novel protocol for persister cells isolation. PLoS One, 2014. 9(2): p. e88660.

18. Germain, E., et al., Molecular Mechanism of Bacterial Persistence by HipA. Molecular Cell, 2013. 52(2):p. 248–254.

19. Semanjski, M., et al., The kinases HipA and HipA7 phosphorylate different substrate pools in Escherichia coli to promote multidrug tolerance. Sci Signal, 2018.11(547).

20. Kaspy, I., et al., HipA-mediated antibiotic persistence via phosphorylation of the glutamyl-tRNA-synthetase. Nature Communications, 2013. 4: p. 3001.

21. Correia, F.F., et al., Kinase Activity of Overexpressed HipA Is Required for Growth Arrest and Multidrug Tolerance in Escherichia coli. Journal of Bacteriology, 2006.188(24): p. 8360–8367.

22. Korch, S.B. and T.M. Hill, Ectopic Overexpression of Wild-Type and Mutant hipA Genes in Escherichia coli: Effects on Macromolecular Synthesis and Persister Formation. Journal of Bacteriology, 2006.188(11): p. 3826–3836.

23. Schumacher, M.A., et al., HipBA-promoter structures reveal the basis of heritable multidrug tolerance. Nature, 2015. 524:p. 59.

24. Schwanhäusser, B., et al., Global quantification of mammalian gene expression control. Nature, 2011. 473:p. 337.

25. Doherty, M.K., et al., Turnover of the Human Proteome: Determination of Protein Intracellular Stability by Dynamic SILAC. Journal of Proteome Research, 2009. 8(1): p. 104–112.

26. Chua, S.L., et al., Selective labelling and eradication of antibiotic-tolerant bacterial populations in Pseudomonas aeruginosa biofilms. Nat Commun, 2016. 7:p. 10750.

27. Spanka, D.T., et al., High-Throughput Proteomics Identifies Proteins With Importance to Postantibiotic Recovery in Depolarized Persister Cells. Front Microbiol, 2019. 10:p. 378.

28. Sulaiman, J.E., C. Hao, and H. Lam, Specific Enrichment and Proteomics Analysis of Escherichia coli Persisters from Rifampin Pretreatment. J Proteome Res, 2018. 17(11): p. 3984–3996.

29. Pu, Y., et al., Enhanced Efflux Activity Facilitates Drug Tolerance in Dormant Bacterial Cells. Mol Cell, 2016. 62(2):p. 284–294.

30. Hansen, S., K. Lewis, and M. Vulić, Role of Global Regulators and Nucleotide Metabolism in Antibiotic Tolerance in Escherichia coli. Antimicrobial Agents and Chemotherapy, 2008. 52(8):p. 2718–2726.

31. Keren, I., et al., Specialized Persister Cells and the Mechanism of Multidrug Tolerance in Escherichia coli. Journal of Bacteriology, 2004.186(24): p. 8172–8180.

32. Prossliner, T., et al., Ribosome Hibernation. Annu Rev Genet, 2018. 52:p. 321–348.

33. Gohara, D.W. and M.F. Yap, Survival of the drowsiest: the hibernating 100S ribosome in bacterial stress management. Curr Genet, 2018. 64(4): p. 753–760.

34. Yoshida, H. and A. Wada, The 100S ribosome: ribosomal hibernation induced by stress. Wiley Interdiscip Rev RNA, 2014. 5(5): p. 723–32.

35. Zhang, Y., et al., HflX is a ribosome-splitting factor rescuing stalled ribosomes under stress conditions. Nat Struct Mol Biol, 2015. 22(11):p. 906–13.

36. Spat, P., et al., Chlorosis as a Developmental Program in Cyanobacteria: The Proteomic Fundament for Survival and Awakening. Mol Cell Proteomics, 2018. 17(9):p. 1650–1669.

37. Varik, V., et al., Composition of the outgrowth medium modulates wake-up kinetics and ampicillin sensitivity of stringent and relaxed Escherichia coli. Scientific Reports, 2016. 6:p. 22308.

38. Denby, K.J., et al., The mechanism of a formaldehyde-sensing transcriptional regulator. Scientific Reports, 2016. 6:p. 38879.

39. Gottesman, S., et al., The ClpXP and ClpAP proteases degrade proteins with carboxy-terminal peptide tails added by the SsrA-tagging system. Genes Dev, 1998.12(9): p. 1338–47.

40. Battesti, A., N. Majdalani, and S. Gottesman, The RpoS-mediated general stress response in Escherichia coli. Annu Rev Microbiol, 2011. 65:p. 189–213.

41. Maciąg, A., et al., In vitro transcription profiling of the σ S subunit of bacterial RNA polymerase: re-definition of the σ S regulon and identification of σ S -specific promoter sequence elements. Nucleic Acids Research, 2011. 39(13): p. 5338–5355.

42. Miura, K., et al., The effects of unsaturated fatty acids, oxidizing agents and Michael reaction acceptors on the induction of N-ethylmaleimide reductase in Escherichia coli: possible application for drug design of chemoprotectors. Methods and findings in experimental and clinical pharmacology, 1997.19(3): p. 147–151.

43. Umezawa, Y., et al., The Uncharacterized Transcription Factor YdhM Is the Regulator of the nemA Gene, Encoding N-Ethylmaleimide Reductase. Journal of Bacteriology, 2008. 190(17): p. 5890–5897.

44. Lilleorg, S., et al., Bacterial ribosome heterogeneity: Changes in ribosomal protein composition during transition into stationary growth phase. Biochimie, 2019. 156:p. 169–180.

45. Keren, I., et al., Characterization and transcriptome analysis of Mycobacterium tuberculosis persisters. mBio, 2011. 2(3): p. e00100–11.

46. Helaine, S., et al., Internalization of Salmonella by Macrophages Induces Formation of Nonreplicating Persisters. Science, 2014. 343(6167): p. 204–208.

47. Ayrapetyan, M., T.C. Williams, and J.D. Oliver, Bridging the gap between viable but non-culturable and antibiotic persistent bacteria. Trends Microbiol, 2015. 23(1):p. 7–13.

48. Song, S. and T.K. Wood, ‘Viable but non-culturable cells’ are dead. Environmental Microbiology, 2021.

49. Cañas-Duarte, S.J., S. Restrepo, and J.M. Pedraza, Novel Protocol for Persister Cells Isolation. PLOS ONE, 2014. 9(2): p. e88660.

50. Shah, D., et al., Persisters: a distinct physiological state of E. coli. BMC Microbiology, 2006. 6(1): p. 53.

51. Dieterich, D.C., et al., Labeling, detection and identification of newly synthesized proteomes with bioorthogonal non-canonical amino-acid tagging. Nature Protocols, 2007. 2:p. 532.

52. Weinert, B.T., et al., Accurate Quantification of Site-specific Acetylation Stoichiometry Reveals the Impact of Sirtuin Deacetylase CobB on the E. coli Acetylome. Mol Cell Proteomics, 2017. 16(5):p. 759–769.

53. Ringquist, S., et al., Translation initiation in Escherichia coli: sequences within the ribosome-binding site. Molecular Microbiology, 1992. 6(9): p. 1219–1229.

54. Baba, T., et al., Construction of Escherichia coli K-12 in-frame, single-gene knockout mutants: the Keio collection. Mol Syst Biol, 2006. 2:p. 2006 0008.

55. Gotfredsen, M. and K. Gerdes, The Escherichia coli relBE genes belong to a new toxin-antitoxin gene family. Molecular Microbiology, 1998. 29(4):p. 1065–1076.

56. Guyer, M., et al. Identification of a sex-factor-affinity site in E. coli as γδ. in Cold Spring Harbor symposia on quantitative biology. 1981. Cold Spring Harbor Laboratory Press.

57. Guzman, L.M., et al., Tight regulation, modulation, and high-level expression by vectors containing the arabinose PBAD promoter. Journal of Bacteriology, 1995. 177(14): p. 4121–4130.

58. Conte, E., et al., pGOODs: new plasmids for the co-expression of proteins in Escherichia coli. Biotechnol Lett, 2011. 33(9):p. 1815–21.

59. Rappsilber, J., M. Mann, and Y. Ishihama, Protocol for micro-purification, enrichment, prefractionation and storage of peptides for proteomics using StageTips. Nature Protocols, 2007. 2:p. 1896.

60. Cox, J. and M. Mann, MaxQuant enables high peptide identification rates, individualized p.p.b.-range mass accuracies and proteome-wide protein quantification. Nature Biotechnology, 2008. 26:p. 1367.

61. Elias, J.E. and S.P. Gygi, Target-decoy search strategy for increased confidence in large-scale protein identifications by mass spectrometry. Nature Methods, 2007. 4: p. 207.

62. Tyanova, S., et al., The Perseus computational platform for comprehensive analysis of (prote)omics data. Nature Methods, 2016.13:p. 731.

63. Huang, D.W., B.T. Sherman, and R.A. Lempicki, Systematic and integrative analysis of large gene lists using DAVID bioinformatics resources. Nature Protocols, 2008. 4:p. 44.

64. Vizcaíno, J.A., et al., 2016 update of the PRIDE database and its related tools. Nucleic Acids Research, 2016. 44(D1):p. D447–D456.

